# CsMT: a robust and streamlined CryoSPARC workflow for cryo-EM reconstruction of microtubules

**DOI:** 10.64898/2026.07.31.741890

**Authors:** Tina Alagha, Asuva Arin, Nicholas Vangos, Helena Goodey-Parfitt, Hai Nguyen Ngo, Nhat Nam Dau, Minh Hoa Nguyen, Thibault Legal, Michael Cianfrocco, Khanh Huy Bui

## Abstract

Microtubules are cytoskeletal filaments that are involved in intracellular transport, cell division, and motility. Despite their biological importance, determining their high-resolution structures via cryo-electron microscopy remains a significant technical challenge due to their polymorphisms and pseudo-helical assembly. Current processing workflows are complex, often requiring the integration of multiple software packages and custom scripts, which creates a steep learning curve for many research groups.

To address these limitations, we introduce CsMT, a streamlined workflow implemented entirely within the CryoSPARC environment and using synthetic references. CsMT simplifies microtubule reconstruction by utilizing a novel protofilament-pair classification approach, which effectively handles the inherent pseudo-symmetry and structural heterogeneity of microtubules with minimal manual intervention. Our workflow is versatile, capable of processing both undecorated and decorated microtubules while accurately determining seams and performing high-resolution refinement.

We demonstrate the efficacy of this workflow by achieving a 2.3 and 2.7 Å resolution reconstruction of homotypic and heterotypic maps of undecorated microtubules, matching the best-resolved microtubule structures in the field. By unifying the pipeline into a single and portable workflow, CsMT enhances reproducibility and accessibility, empowering more laboratories to explore the structural biology of microtubules and associated proteins, yielding new insights into their function.

## Introduction

Microtubules (MTs) are dynamic and cylindrical cytoskeletal polymers of α-β tubulin heterodimers that polymerize longitudinally to form protofilaments (PFs). These structures support a wide variety of cellular functions such as cellular structure, cell division, and intracellular transport. MTs are highly dynamic polymers that undergo stochastic cycles of growth and shrinkage, a process known as dynamic instability. MTs rely on a diverse array of microtubule-associated proteins (MAPs) to balance stability with disassembly. Beyond regulating polymer dynamics, MAPs govern lattice organization, the formation of higher-order structures, and integration with other cellular components (Akhmanova and Kapitein 2022). High-resolution cryo-electron microscopy (cryo-EM) studies of reconstituted MTs in complex with MAPs are essential for understanding how these proteins regulate MT organization and function. However, MTs remain challenging structural targets because they adopt pseudo-helical symmetries due to a notable lattice discontinuity - the “seam”. Microtubule seams exhibit lateral contacts that differ from the canonical B-lattice homotypic interactions; instead, they form A-lattice heterotypic interactions that break helical equivalence and create a structurally distinct interface along the MT.

Identifying the seam is essential for asymmetric reconstruction and the differentiation of α-and β-tubulin. Although the two subunits are structurally similar, β-tubulin is distinguished by a longer M-loop and C-terminal tail (Nogales et al. 1999). Unless the seams of the MTs are identified and aligned correctly during reconstruction, forcing symmetry onto pseudo-helical polymers can obscure such structural details (Cope et al. 2010). While MTs in the cell are found mostly containing 13-PFs, MTs ranging from 9-PF to 16-PFs are found by reconstitution *in vitro* (Pierson, Burton, and Himes 1978; Chrétien and Wade 1991). The ratio of different types of MTs depends on nucleotide and ligand used for polymerization (Sui and Downing 2010) and can be influenced by MAP during co-polymerization (Bechstedt and Brouhard 2012). MTs can vary in lattice spacing, or the distance between α and β tubulin dimers, as GTP-bound tubulin has a 2.3 ± 0.2 % wider spacing than GDP-bound tubulin (Hyman et al. 1995; Alushin et al. 2014). When a MT has 13-PFs, the PFs run almost parallel to the axis of the MT. Due to geometric constraints, deviation from 13-PF MT causes a protofilament tilt relative to the main longitudinal axis of the MT, i.e. a skew angle (Fig. 1A). In addition, MAPs can bind at different MT interfaces such as on β-tubulin like kinesin (Kikkawa et al. 1995) between four tubulin dimer like End binding protein (Zhang et al. 2015), or at the seam like SPEF1 (Legal et al. 2025). Therefore, MTs are inherently variable structures, differing in PF number, lattice spacing, super twisting and MAP decoration. These factors together complicate cryo-EM reconstruction workflows.

**Fig. 1:**
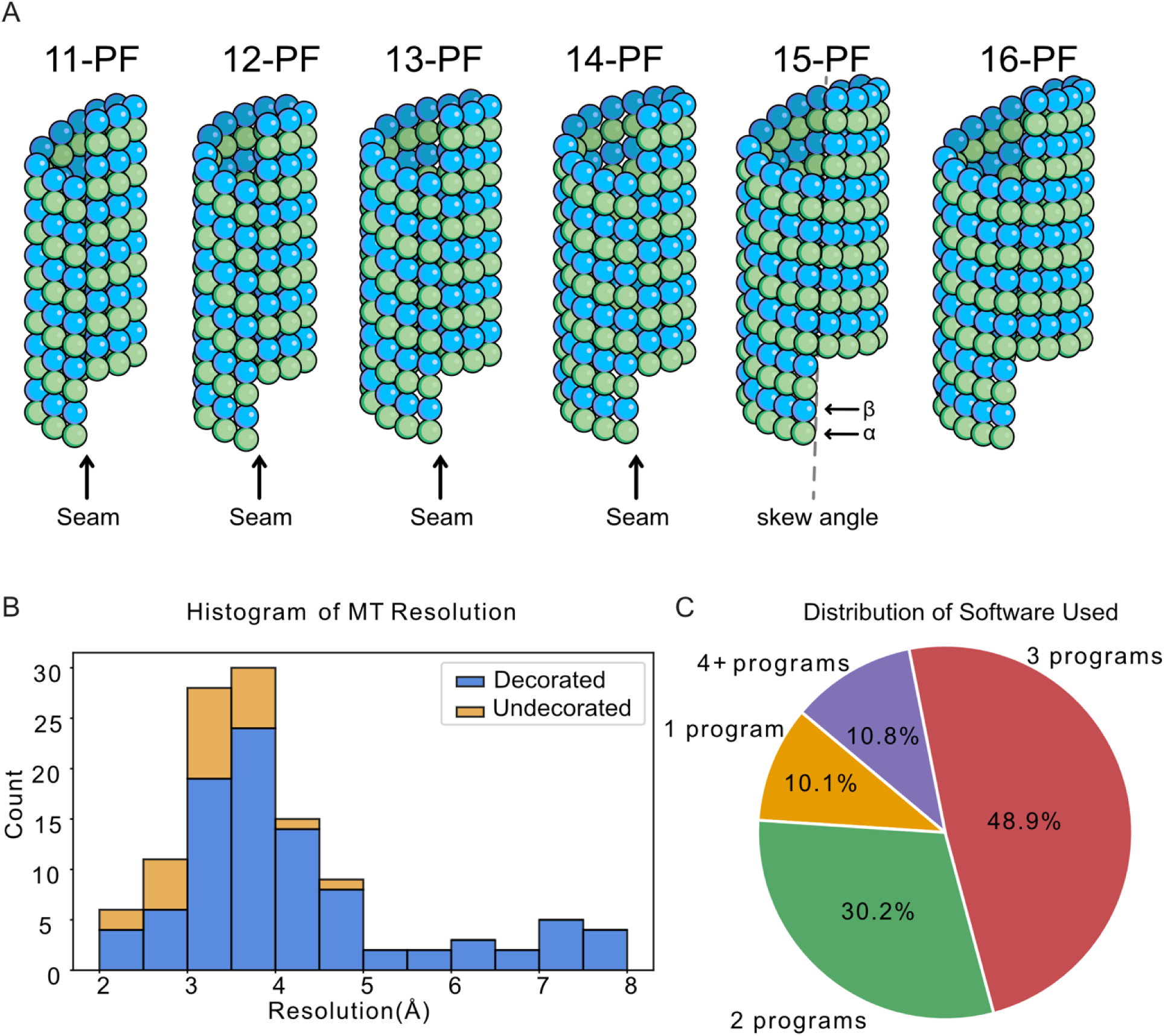
Overview of MT structures & cryo-EM reconstruction statistics. A. MT polymorphism with different PF numbers. B. Histogram of resolution of Cryo-EM maps of decorated and undecorated MTs deposited in Electron Microscopy Databank as of 13^th^ July 2026. C. Software used for MT analysis.

The truly helical MT forms such as 15-and 16-PF MTs can be solved by cryo-EM using classical Fourier-based helical reconstruction methods (Hirose et al. 1997; Kikkawa et al. 1995; Sosa et al. 1997) and the Iterative Helical Real Space Reconstruction (IHRSR) approach (Egelman 2000). With the advances of cryo-EM and the flexible nature of biological filaments, the IHRSR approach has become the standard approach to solve helical biological structure (He and Scheres 2017). For pseudo-helical MTs, different approaches have been developed, which can be divided into two broad categories: (i) seam search-based (Sindelar and Downing 2007; Zhang and Nogales 2015; Zhang et al. 2026; Cook et al. 2020) and (ii) PF-classification based methods (Debs et al. 2020; Vangos et al. 2026).

The principle of the seam-search is to find the seam of the MT based on the assumption that there is only one continuous seam along the same MT. This is achieved through three-dimensional (3D) alignment and classification and smoothing of the phi alignment angles to find the continuous seam (Zhang and Nogales 2015; Zhang et al. 2026; Cook et al. 2020). In contrast, the PF-classification based method does not assume a single seam but focuses on classification on a single or pair of PF to distinguish the 4-nm shift classes of α-and β-tubulins (Vangos et al. 2026; Debs et al. 2020). In both approaches, focus refinement on only one or two PFs rather than the entire MT (Vangos et al. 2026; Zhang et al. 2026; Debs et al. 2020; Luo et al. 2026) yields superior resolution by mitigating the effects of MT flexibility and lattice distortions introduced during vitrification. Consequently, MT structure resolution has steadily improved over the years, recently reaching beyond 2.5 Å (Fig. 1B). Both these approaches work on MAP-decorated MTs but also work on undecorated MTs with some modifications (Cook et al. 2020). However, structure determination of undecorated MTs is more challenging than that of decorated MTs due to the high structural similarity between α-and β-tubulins.

With these approaches, the cryo-EM workflow for MT tends to use multiple programs and scripts such as Frealign (Grigorieff 2016), RELION (Scheres 2012) and CryoSPARC (Punjani et al. 2017) (Fig. 1C). In particular, seam-search pipelines often involve manual picking, external scripts, and particle pre-averaging (Zhang and Nogales 2015; Zhang et al. 2026; Cook et al. 2020). This complexity poses a barrier for new cryo-EM users and laboratories lacking specialized expertise in this area. As a result, routine MT reconstruction remains less accessible, limiting the diversity of MAP-MT complexes and MT states that can be systematically characterized. Single package approaches have been proposed using the two most popular single particle cryo-EM packages: RELION (Vangos et al. 2026) and CryoSPARC (Zhang et al. 2026). Recently, MiCSPARC is developed as a new CryoSPARC pipeline utilizing CryoSPARC in combination with external scripts for fitting, angle unification and seam search. CryoSPARC offers better computational optimization for large datasets. Furthermore, CryoSPARC has Workflow feature, which can be generated, exported and shared between different laboratories, allowing consistent analysis and ease-of-use for new users. However, this feature cannot be implemented easily with the presence of external script.

Here, we introduce CsMT, a PF-classification-based pipeline to solve the accessibility and reproducibility of MT reconstruction. CsMT is a minimal CryoSPARC-only workflow utilizing a novel PF-pair classification approach that does not require any external software or custom scripts. CsMT is designed to reconstruct both decorated and undecorated MTs to high resolution. By providing a shareable and easily adjustable CryoSPARC workflow, CsMT delivers consistent, high-resolution results with minimal manual effort, making it a more reliable and efficient alternative to existing approaches.

## Results

### CsMT can reconstruct undecorated MT using a novel PF-pair classification approach

The CsMT pipeline is a PF-classification approach with similarity to a method developed using RELION (Vangos et al. 2026). For single PF reconstruction, our method is essentially identical to the novel method implemented in RELION (Vangos et al. 2026). For our CryoSPARC workflow, we introduced two important distinctions: (i) 4-nm filament picking instead of 8-nm filament picking to avoid particle duplication; and (ii) a robust “PF-pair classification” logic, allowing us to reconstruct two-PF seam and non-seam structures without the need of any external scripts. The full workflow is detailed in Fig. 2.

**Fig. 2:**
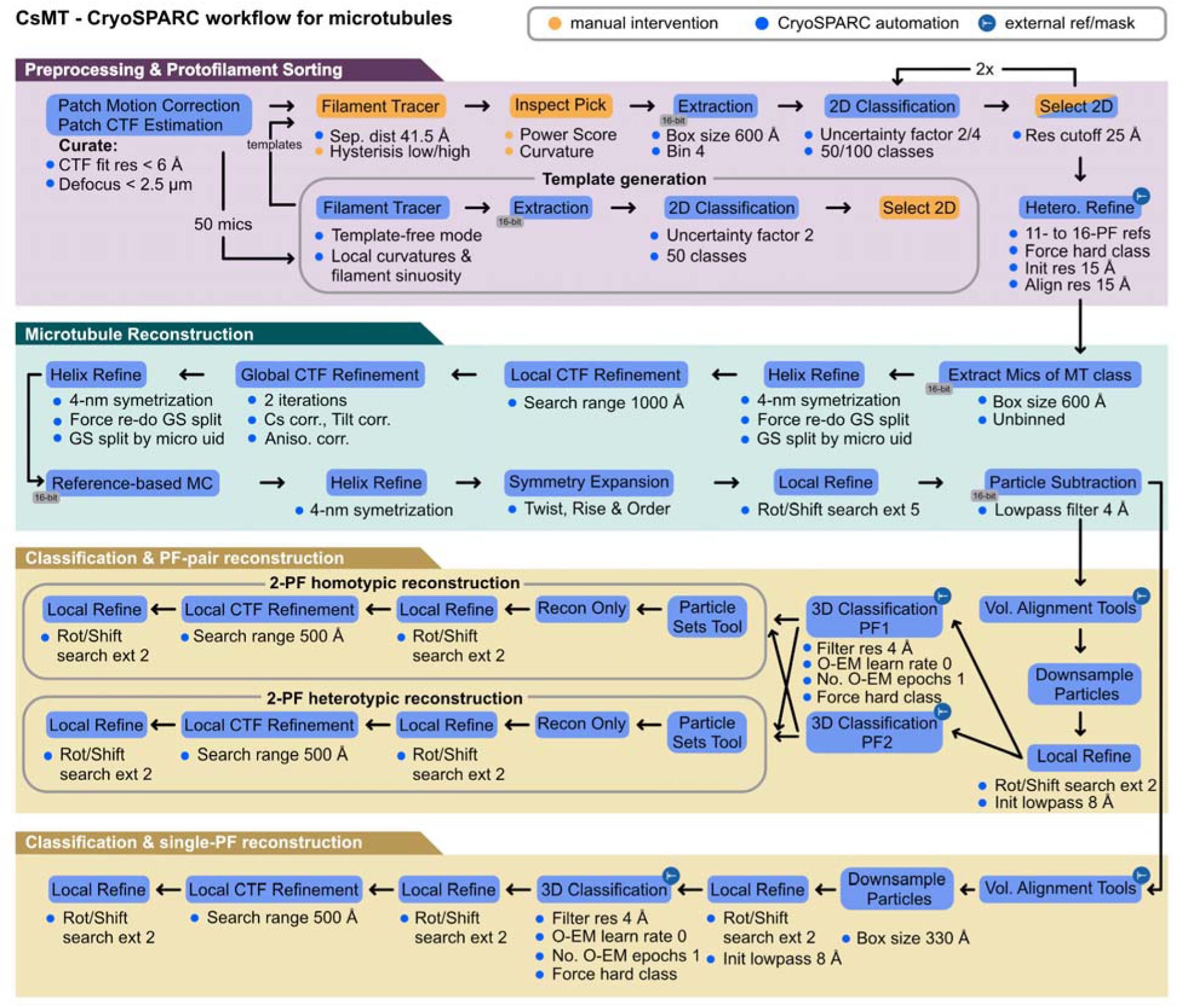
**CsMT processing workflow.**

### Preprocessing

Beam-induced motion and contrast transfer functions (CTF) were corrected using Patch Motion Correction and Patch CTF Estimation. Micrographs were discarded if the CTF fit resolution exceeded 6 Å or the defocus exceeded 2.5 µm.

### Filament Picking

MTs were picked in two rounds using the Filament Tracer. First, a template-free picker was run on 40–100 representative micrographs, and the resulting particles were 2D-classified to generate initial templates. To ensure rare polymorphs (e.g., 12-PF MTs) are captured, more micrographs should be used for this initial template generation. These templates were then deployed across the entire dataset with a 41 Å picking interval to sample both α-and β-tubulin subunits indiscriminately. Finally, two rounds of 2D classification were performed to remove false positives and damaged filaments.

### Classification and Helical Refinement

MT polymorphism was resolved via heterogeneous refinement in CryoSPARC using synthetic 11-to 16-PF references. While 11-and 16-PF structures were rare and can be omitted, including 12-to 15-PF references is essential for clean separation. Particles from the most abundant class (typically 13-or 14-PF) were selected, recentered, and re-extracted unbinned. Initial helical refinement was performed, yielding a 4-nm repeat reconstruction with averaged α-/β-tubulin density. To maximize resolution, helical refinements were iteratively interleaved with Local CTF Refinement, Global CTF Refinement, and Reference-based Motion Correction.

### Single-PF Refinement

To resolve the α-/β-tubulin registry, a signal subtraction mask was applied to isolate a single PF (Fig. 3). Subtracted particles were recentered, cropped, and subjected to Local Refinement. Next, 3D classification with two synthetic references— representing "α-first" and "β-first" orientations—separated the particles into two 4-nm axially shifted classes (typically a ∼50:50 distribution). Local refinement of either class resolves the correct 8-nm tubulin dimer repeat. Final maps were further improved by optimizing per-particle defocus values via Local CTF Refinement, followed by a final Local Refinement.

**Fig 3:**
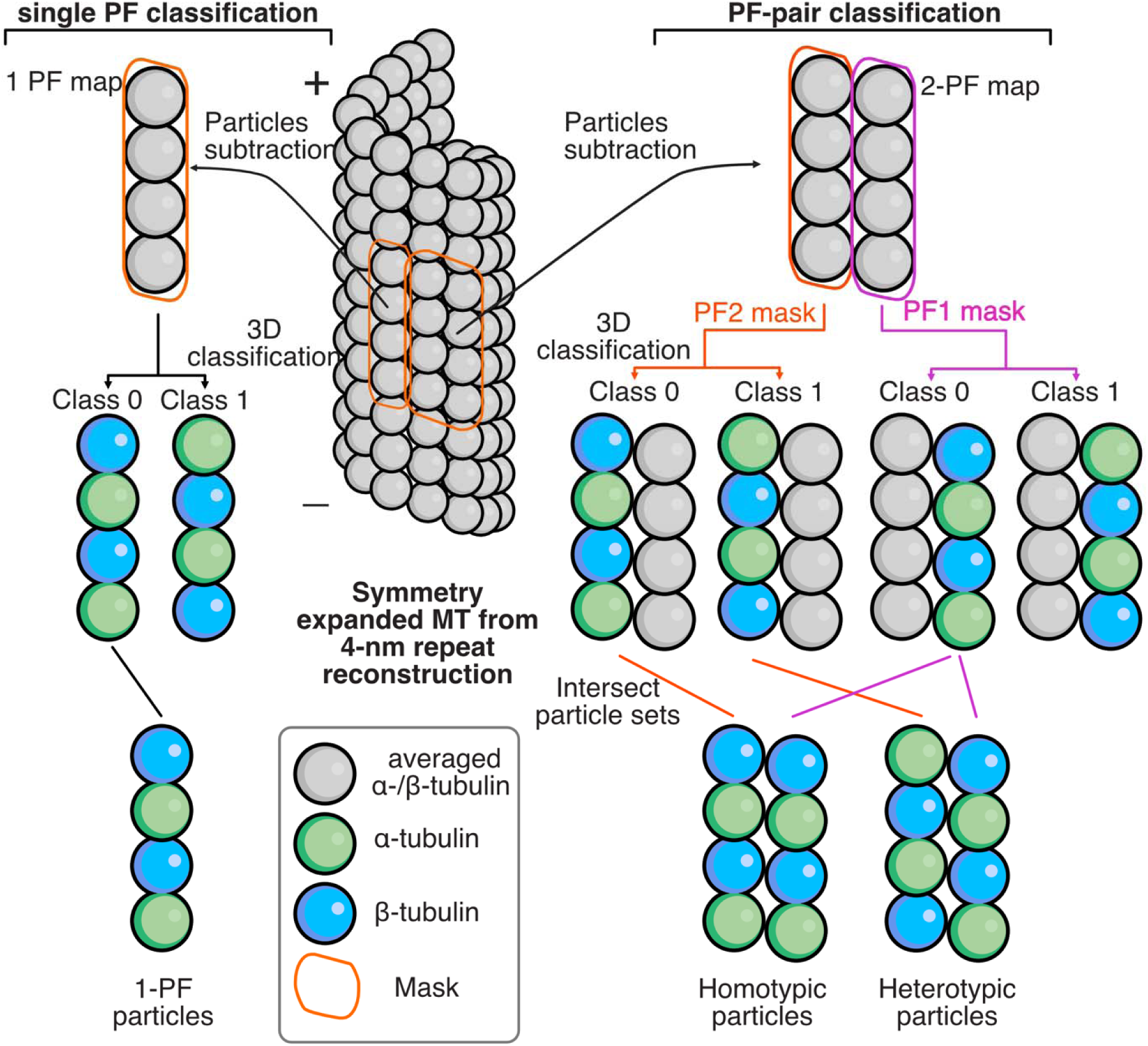
**Principle of classification and grouping for 1-PF and 2-PF pair reconstruction.**

### Two-PF Refinement

To visualize both homotypic (B-lattice) and heterotypic (seam) lateral interactions, a two-PF refinement approach was developed. Density corresponding to a block of 3 x 2 tubulin dimers across two adjacent PFs was isolated via signal subtraction (Fig. 3). Subtracted particles were recentered, cropped, and subjected to Local Refinement. To determine the α-/β-tubulin registry for each filament, focused 3D classification was performed independently for each PF using 2-PF masks containing two dimers each (PF1 and PF2) and synthetic references representing the 4-nm axial shift.

Intersecting these classification results isolated specific lattice architectures: particles belonging to discordant classes (e.g., class 0 for PF1 and class 1 for PF2) represented the heterotypic seam, whereas concordant classes (class 0 for both PFs) represented the homotypic B-lattice. These separated populations were then individually optimized using a cycle of Local Refinement, Local CTF Refinement, and a final Local Refinement.

### Map Post-Processing

Final reconstructions were post-processed using Phenix density modification (Terwilliger et al. 2020) or EMReady2 (Cao et al. 2026) to improve interpretability for downstream atomic modeling.

Applying this workflow to the cryo-EM data of taxol stabilized human MTs (Vangos et al. 2026), we obtain a 2.3 Å resolution for single PF reconstructions, and 2.3 Å and 2.7 Å resolution 2-PF maps of homotypic and heterotypic, respectively (Fig. 4A-C and Fig. S1). The resolution of the maps are comparable to the structures obtained by a novel PF-classification based RELION workflow (Vangos et al. 2026).

**Fig. 4:**
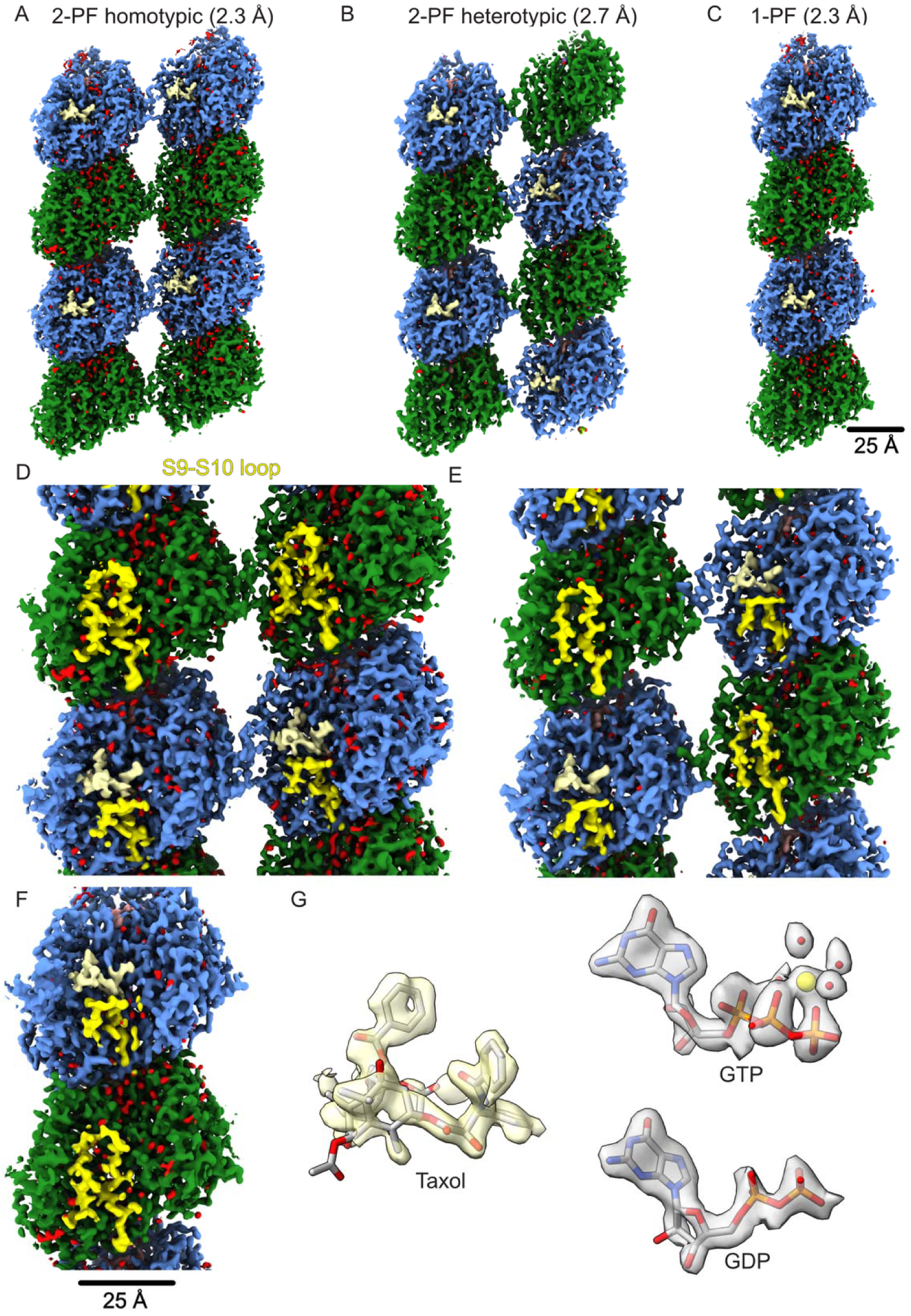
Processing of undecorated taxol bound human MTs using CsMT workflow. (A) 2-PF homotypic (B-lattice) map at 2.3 Å resolution (2.2 Å using density modification, EMD-78103). (B) 2-PF heterotypic (A-lattice) map at 2.7 Å resolution (2.6 Å using density modification, EMD-78104). (C) 1-PF map at 2.3 Å resolution (2.2 Å using density modification, EMD-78112). Maps displayed were post-processed using Phenix density modification. Color: α-tubulin: green; β-tubulin: light blue; Taxol: beige; Water: red; (D-F) Magnified views of 2-PF homotypic (D), 2-PF heterotypic (E) and 1-PF (F) maps show distinct densities of S9-S10 loops, a feature that distinguishes α*-*and β-tubulin. (G) Densities of taxol, GDP and GTP from 1-PF map. The GTP density shows magnesium (yellow) and water densities (red).

When running our map through density modification (Terwilliger et al. 2020), the resolutions for our single-PF and 2-PF map are improved to 2.2 Å. All our map shows clearly the S9-S10 loop and density in the GTP & GDP and taxol of α-tubulin and β-tubulin (Fig. 4D, E, F). As testament for our high-resolution map, we can visualize the taxol density in our map with unprecedented clarity (Fig. 4G).

Our 2-PF seam map is the best resolved seam reported so far (Fig. 4E). This is a clear indication that our method works well to resolve the seam structure. Using intersection action from Particle Set Tools from CryoSPARC in combination with our novel PF-pair classification, we can effortlessly map particle uid for seam reconstruction without any custom scripts for seam search or seam mapping.

### CsMT can flexibly accommodate decorated/MAP-bound MTs

Though our workflow is designed for undecorated MTs, simple modifications can be done to accommodate either fully or partially decorated MTs. For fully decorated MTs, we can perform 3D classification to obtain the correct α-/β-tubulin registry at the same time with the MAP classification using a 4-nm repeating mask containing also the MAP densities (Fig. 5A). For partially decorated MTs, we should process 3D classification for the correct α-/β-tubulin registry like the undecorated workflow to obtain the correct α-/β-tubulin registry map. Then we can use a mask for 3D focus classification on the MAP density to separate decorated and non-decorated particles (Fig. 5B).

**Fig. 5:**
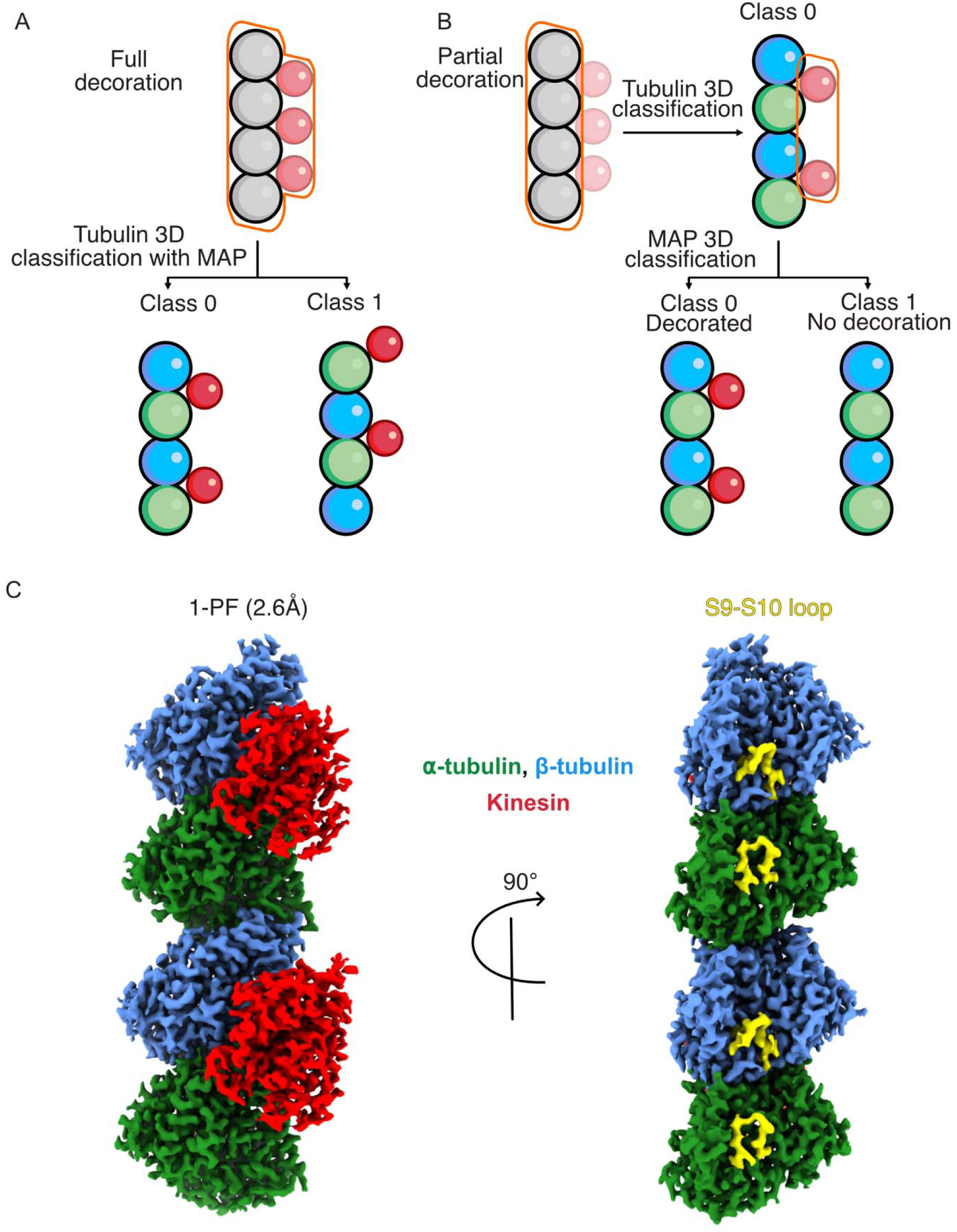
CsMT workflow for decorated MT. (A) If the MT is fully decorated, the tubulin classification step in CsMT from the α-/β-averaged map can also include the decorated MAP by making the classification mask containing also the 4-nm averaged MAP density. (B) If the MT is partially decorated, the tubulin classification step in CsMT is done strictly based on the tubulin mask and the synthetic references of α-first and β-first PF. After that, the correctly 8-nm registered PF map is subjected to another classification to separate the decorated PF particles from the undecorated PF particles. (C) PF map of tubulin decorated with KIF5A, EMD-78245 reconstructed from EMPIAR-13372 (Zhang et al. 2026) at 2.6 Å resolution.

We applied our workflow on data of MTs decorated with kinesin head domains (EMPIAR-13372) (Zhang et al. 2026). We obtained a map at 2.6 Å resolution where it clearly shows the α-/β-tubulin and the kinesin density (Fig 5C, Fig. S2). Our resolution is slightly better than the reported resolution using MiCSPARC of 2.8 Å (Zhang et al. 2026). This demonstrates that the CsMT workflow is also efficient for decorated MTs.

### CsMT workflow allows optimization for high-resolution reconstruction

With our efficient CsMT workflow and current throughput of data collection (5000-10000 movies per day), we expect that we can routinely achieve ∼2.5 Å resolution. To obtain high-resolution reconstruction of MTs, there are various factors for consideration. Since the CsMT workflow is very simple to implement, we were able to test various parameters to choose the optimum for our data in terms of high resolution and processing time.

### Defocus and Box Size Optimization

High-defocus data inherently limits high-resolution reconstruction due to CTF signal delocalization. For a 300 kV microscope and a 600 Å box size, achieving a 2.5 Å target resolution requires limiting defocus to less than 2.2 µm. This restriction constrains spatial delocalization Δ ·*r* =λ Δ*z*/*d* (λ: electron wavelength; · Δ*z* : defocus, : resolution) to a maximum of 175 Å, matching the padding threshold outside the MT walls (Rosenthal and Henderson 2003). Comparable delocalization constraints apply when recentering and cropping the 2-PF particles. Consequently, a conservative defocus threshold of less than 2.0 µm was enforced to maximize high-resolution signal retention.

### Filament Picking Optimization

Although most of the pipeline is automated (Fig. 2), the template-based filament picking and inspection stages require careful manual tuning, as optimal parameters vary between datasets. Initial optimization should be performed on a subset of approximately 50 micrographs by systematically testing the low and high hysteresis thresholds in the Filament Tracer job, alongside the Normalized Cross-Correlation power and score thresholds in the Inspect Picks step.

In high magnification dataset such as EMPIAR-13372, there is a significant under pick of particles due to short filaments and large box size required. In this case, a filament extrapolation method such as one introduced by MiCSPARC (Zhang et al. 2026), can nicely improve the particle picking. As shown in this work, spending time to optimize the filament picking can achieve good particle picking coverage. Therefore, external scripts are best reserved for advanced users, as custom scripting interrupts the seamless CryoSPARC workflow.

### Heterorefinement optimization

This step is crucial for the success of the workflow. However, it is not always easy for stably running this step as the minor classes such as 11-PF, 12-PF, 15-PF and 16-PF can be as low as 1% of the particles. This leads to attraction problems when larger and better resolved classes will attract all the particles from other minor classes. For CryoSPARC, restricting the initial resolution to 15 Å and alignment resolution to 15 Å and turning on Force Hard Classification helps significantly with the convergence of the PF sorting. Additionally, limiting the reference framework to 12-PF through 15-PF proves beneficial; the 12-PF classification absorbs neighboring 11-PF particles, whereas the 15-PF class similarly assimilates 16-PF particles. However, the sorting will still struggle with small datasets when the particles from minor classes will not yield a ∼15 Å map. In this case, perhaps limit the references only to the major classes (13-and 14-PF) would be more appropriate.

### Mask Size Optimization

For PF classification, focused masks containing two tubulin dimers per PF (∼400 kDa molecular weight) provided sufficient signal for accurate alignment, performing comparably to larger three-dimer masks (∼600 kDa). Furthermore, during the final 2- PF local refinements, the smaller two-dimer x 2-PF mask consistently yielded marginal improvements in resolution: 2.28 Å versus 2.32 Å for the homotypic B-lattice, and 2.70 Å versus 2.74 Å for the heterotypic seam.

### Centering, Cropping, and Computational Efficiency

Centering and cropping the 2-PF density based on the focused refinement mask is critical for both computational performance and rotational accuracy. Centering places the fulcrum of the rotational search directly at the center of mass of the focused mask, improving alignment precision. Structurally, cropping the particles from a 720 x 720-pixel box down to a 400 x 400-pixel box reduced the storage footprint of a 2.6-million particle dataset from ∼2.4 TB to ∼770 GB (a 68% reduction). This significantly optimized data management; for a subset of 1 million particles, SSD caching took only 2 hours over our network, and the subsequent local refinement was completed in 4 hours on a single NVIDIA RTX 3090 GPU.

### Classification Strategy

Unlike RELION, CryoSPARC does not feature a regularization parameter (T-value) to tune 3D classification sensitivity. To accurately resolve the expected ∼50:50 distribution between the "α-first" and "β-first" classes, specific classification parameters must be enforced. Activating *Force Hard Classification* prevents particles from weights-blending across classes, while setting the *O-EM learning rate* to 0 and enforcing *Filter to Resolution* at 4 Å ensures the initial synthetic references strictly guide the Full-EM classification phase without drifting.

### Gold-Standard Fourier Shell Correlation (FSC) Validation

Due to particle overlap along the helical axis, filament reconstructions are highly susceptible to resolution inflation caused by duplicate or overlapping particles leaking into both half-datasets during refinement. This explains the fact that certain MTs maps appear worse than the reported resolution. To prevent this, particles belonging to the same physical filament must be restricted to the same half-set for Gold-Standard processing.

While maintaining filament IDs across software suites can be problematic due to data loss during conversion, our workflow mitigates this by enforcing a 4-nm picking interval and 4-nm helical symmetry during refinement. However, because filament-ID-based splitting can still be imperfect, partitioning the gold-standard split strictly by micrograph ID (a feature native to CryoSPARC) provides the most rigorous and accurate method for FSC resolution estimation in large datasets.

## Discussion

In this work, we developed CsMT, a CryoSPARC workflow to reconstruct MTs. Notably, our new processing workflow, available as import-ready.json files (https://github.com/builab/CsMT), achieved 2.3 Å global resolution for single-PF map of MTs in approximately 157 hours using one GPU processing of ∼3,200 movies. Compared to the seam search approach, our approach does not depend on picking a long filament. Thus, we do not need either manual picking (Cook et al. 2020) or scripts for filament interpolation (Zhang et al. 2026) or seam smoothing after classification.

Furthermore, our analysis suggests that there is not a single seam along the MT. The number of particles of seam vs. B-lattice is about 1 to 4 for our 13-PF MTs, instead of 1:13 ratio if a single seam assumption is true. Our observation is supported by good resolution obtained from our seam map, which suggests the majority of the classification is correct as otherwise, 50% of the particles would be assigned wrongly in our seam particle set based on one seam assumption. A cryo-electron tomography study using kinesin-decorated MTs visualizes multiple seams (Guyomar et al. 2022) which supports our observation. On the other hand, the seam search approach works quite well, therefore in vitro reconstituted MTs likely have a main seam running along but might contain other seams due to imperfect polymerization.

The single-PF approach likely yields better resolution than the 2-PF approach, as it is less impacted by biological flexibility and the flattening of MTs during vitrification. On the other hand, the PF-pair approach allows a nice visualization of the seam or the B-lattice instead of a single PF. This is probably a big advantage if the MAP of interest is binding between two PFs such as EB proteins binding to between four tubulin dimers (Maurer et al. 2012) or SPEF1 binding between the seam (Legal et al. 2025).

The seam search approach allows the reconstruction of the full MT (Zhang and Nogales 2015; Zhang et al. 2026). However, reconstruction of the full MT can be achieved efficiently by combining the seam using multiple copies of the 2-PF maps in ChimeraX using ‘volume maximum’ command (Legal et al. 2025). Furthermore, it is also possible to use scripts to analyse distorted MTs or lattice defects such as demonstrated previously (Debs et al. 2020).

Towards the future, a single workflow CsMT might be able to be tweaked for fully autonomous analysis. All the references and masks can be re-used if we use the same magnification. Selection of 2D classes can be done automatically in a large dataset with a resolution threshold or with a tool like CryoSIFT (Schäfer et al. 2025). The only step that requires manual optimization is filament picking and inspect picking. In a large dataset, it is possible to use stringent criteria, and we are still able to pick enough particles for reconstruction.

As cryo-EM technology continues to evolve, achieving sub-2 Å resolution for MTs is becoming increasingly feasible using advanced detectors and larger datasets. Our CsMT framework contributes to the standardization of the MT reconstruction process, enabling researchers to prioritize sample optimization and data quality to better elucidate MT structure and function.

## Methods

### EMDB Map Statistics

To query the MT maps in the EMDB, we used the following query “title:”microtubule" NOT structure_determination_method:"singleParticle"”.

The resolution histogram was done using a Python script with bin of 0.5 Å resolution with a resolution cutoff of 8 Å.

For the software used statistics, the software reported in EMDB are not accurate. Therefore, we read all the manuscripts and annotated the software manually. The plot was then plotted using Python script.

### Processing of undecorated MT dataset

The dataset of taxol stabilized human MTs was used previously (Vangos et al. 2026).

All data processing was executed within CryoSPARC v5.0.2 utilizing a cluster node equipped with two NVIDIA RTX 3090 GPUs. The raw dataset consisting of 3,106 movie stacks was subjected to patch motion correction and patch CTF estimation. Micrograph curation was performed to discard low-quality images, retaining only those with a CTF fit resolution better than 6.0 Å and a defocus value below 2.5 µm.

An initial particle picking round was performed on a representative subset of 300 micrographs using the Filament Tracer utility, configuring a characteristic filament diameter of 220 Å and a periodic unit length of 41.5 Å. This generated an initial stack of 25,608 particles, which were subjected to 2D classification into 50 classes with an initial uncertainty factor of 2. Four well-resolved, representative MT classes were selected to serve as templates for comprehensive picker targeting. The comprehensive Filament Tracer run was then deployed across the curated micrograph pool utilizing the selected templates with a 220 Å diameter, 41.5 Å pitch, low/high hysteresis thresholds of 91 and 98, and a standard Gaussian blur factor of 0.2. Following an Inspect Picks step to optimize the power score and filament curvature thresholds, a total of 320,023 particles were selected. Particles were extracted using a 720-pixel box and Fourier-cropped to 180 pixels. The particle stack was cleaned via two successive rounds of 2D classification into 50 and 100 classes, with uncertainty factors set to 2 and 4, respectively, yielding 312,048 clean particles finally.

To resolve structural heterogeneity arising from MT polymorphism, the particle stack was subjected to Heterogeneous Refinement using synthetic 12-, 13-, 14-, and 15-PF MT starting volumes as reference models (Vangos et al. 2026). A subset of 200,251 particles assigned to the 13-PF classes was selected and re-extracted at un-binned pixel size with a 720-pixel box. This stack was processed via Helical Refinement by enforcing 13-fold helical symmetry parameters with a rise of 9.59 Å and a twist of 27.69°, resulting in a consensus reconstruction at 3.2 Å resolution. Structural details were further improved through sequential rounds of Local CTF Refinement and Global CTF Refinement, followed by another round of Helical Refinement, driving the resolution to 3.0 Å. Subsequent Reference-Based Motion Correction paired with a final consensus Helical Refinement yielded a 2.9 Å map. The 13-PF consensus particle stack was then symmetry-expanded based on the refined helical operators, yielding 2,600,416 symmetry-expanded units. Following a Local Refinement step that reached 2.8 Å, a strict orientation filter was applied to remove all particles with a tilt angle greater than 20°. The remaining particles were prepared for focused classification by performing signal subtraction targeting either individual PF (1-PF) or pairs of adjacent PFs (2-PF).

The 2-PF subtracted particle stack was centered using the Volume Align Tool and cropped to a 400-pixel box via Downsampling Particles. An initial Local Refinement of this stack yielded a 2.5 Å reconstruction. To break the pseudo-symmetry and resolve the true 8-nm repeating registry of the α-/β-tubulin heterodimers, targeted 3D classification strategies were executed. Synthetic references containing 3 x 2 tubulin dimers with a defined α-/β-first starting configuration were assembled in UCSF ChimeraX from an AlphaFold model and converted to 4 Å resolved density maps using ChimeraX molmap command. Custom masks for classification were built by mapping the PDB model to a 15 Å density in ChimeraX, followed by processing via CryoSPARC Volume Tools using a 4-pixel dilation radius and a 12-pixel soft padding width. Using distinct PF1 and PF2 masks, two independent 3D classification tracks separated the particles into two categories, achieving an identical 51.3% to 48.7% distribution across both runs.

For the 2-PF homotypic reconstruction, particles belonging to Class 1 from both classification tracks were pooled together to form a subset of 997,466 particles. Initial volume generation was completed using Homogeneous Reconstruct Only, followed by Local Refinement that yielded a 2.5 Å map. Applying a final Local CTF Refinement with a 500 Å search range coupled to localized refinement yielded a final homotypic map at 2.3 Å resolution. For the 2-PF heterotypic reconstruction, particles from Class 0 of the PF1 classification track and Class 1 of the PF2 classification track were combined to isolate the heterotypic assembly, resulting in 265,812 particles. This subset was processed through an identical workflow of Homogeneous Reconstruct Only followed by Local CTF Refinement and Local Refinement, resulting in a final heterotypic map at 2.7 Å resolution. For the 1-PF subtracted track, an identical signal subtraction and downsampling protocol was followed. This subset required only a single classification track to sort the tubulin registry, resulting in a final homogeneous pool of 1,275,502 particles. Final Local Refinement of this dataset produced a high-resolution map at 2.3 Å.

Final unsharpened maps were post-processed using Phenix Density Modification (Terwilliger et al. 2020) to maximize feature contrast and connectivity. Structural visualizations and figures were generated using UCSF ChimeraX (Meng et al. 2023). To facilitate domain coloring and analysis, atomic coordinates of the tubulin PF (Vangos et al. 2026) were rigid-body fitted into the 1-PF, 2-PF homotypic, and 2-PF heterotypic reconstructions.

### Processing of decorated MT dataset

The kinesin-decorated, GMPCPP-stabilized MT dataset was obtained from EMPIAR (EMPIAR-13372) (Zhang et al. 2026). Similar to the undecorated MT dataset, we processed the raw dataset within CryoSPARC v5.0.2 utilizing the same cluster node by tweaking some of the parameters in the workflow.

The raw dataset comprised 1,676 micrographs that subjected to motion-correction and patch CTF estimation. Like the undecorated dataset, micrograph curation was performed to retain only images with a CTF fit resolution better than 6.0 Å and a defocus below 2.5 µm.

An initial particle picking round was performed to generate templates on a representative subset of 50 micrographs using the Filament Tracer utility, configuring a characteristic filament diameter of 360 Å and a periodic unit length of 82 Å, yielding an initial stack of 664 particles.

These were subjected to 2D classification into 10 classes with an initial uncertainty factor of 2, and two well-resolved, representative MT classes with clear kinesin decoration were selected to serve as template for picking across the full dataset. The comprehensive Filament Tracer run used the same 360 Å diameter and 82 Å pitch with low/high hysteresis thresholds of 91 and 98 and a Gaussian blur factor of 0.15. Following an Inspect Picks step to optimize the power score and filament curvature thresholds, a total of 47,167 particles were selected. Particles were extracted with a 938-pixel box and Fourier-cropped to 180 pixels and cleaned via two rounds of 2D classification into 100 classes, with an uncertainty factor set to 4, yielding 24,076 clean particles.

Heterogeneous Refinement was performed using the same references from 12-PF to 15-PF. The 19,888 particles assigned to the 14-PF class were selected for Extraction as this class is 85.58% among all the particles in the dataset. The Extraction was done with a 938-pixel box and 720-pixel Fourier-cropped box. Helical Refinement of the stack processed with a rise of 8.94 Å and a twist of 25.76° produced a consensus reconstruction at 3.6 Å resolution. Sequential rounds of Local and Global CTF Refinement followed by another Helical Refinement improved resolution to 3.1 Å. The 14-PF particle stack was then subjected to Symmetry expansion on the refined helical operators, yielding 553,140 symmetry-expanded particles. A subsequent Local Refinement with a full-coverage mask reached 3.0 Å, with no particle rejection, and the stack was prepared for subtraction targeting 1-PF with kinesin using a subtraction mask.

The subtracted particle stack was centered using the Volume Align Tool and cropped to a 400-pixel box via Downsampling Particles. An initial Local Refinement of this stack yielded a 2.8 Å reconstruction. To define α-/β-tubulin starting in the PF, 3D classification strategies were executed with two synthetic references containing 3 x 2 tubulin dimers with a defined α-/β-first starting configuration were assembled in UCSF ChimeraX from PDB 9T17 (Zhang et al. 2026) and converted to 4 Å resolution density maps using ChimeraX molmap command.

3D classification tracks separated the particles into two categories, achieving a 50.5% to 49.5% distribution. One of the classes is selected for final local refinement, resulting in 273,885 particles. Local CTF Refinement followed by a final Local Refinement is done with initial low pass resolution 8Å using a mask covering two tubulin dimers and kinesins, producing the final map at 2.6 Å.

All structural figures were prepared in UCSF ChimeraX (Meng et al. 2023). To facilitate domain coloring and analysis, atomic coordinates of the tubulin and kinesin from Zhang et al., 2026 (Zhang et al. 2026) were rigid-body fitted into the 1-PF map.

## Data & Software Availability

The CryoSPARC workflow json file is available at: https://github.com/builab/CsMT

Data and scripts to plot MT resolution histogram and software usage is also available in the GitHub page.

EM maps of 2-PF homotypic, 2-PF heterotypic and 1-PF of taxol stabilized human MT, and 1-PF of MT with KIF5A have been deposited in the Electron Microscopy Data Bank (EMDB) under accession numbers: EMD-78103, EMD-78104, EMD-78112, and EMD-78245.

## Acknowledgement

We thank Drs. Michal Wieczorek, Hugo Munoz Hernandez, Pavel Filipcik for sharing the movies of their dataset of MT with KIF5A. K.H.B. is supported by grants from the Canadian Institutes of Health Research (PJT-190195) and the Natural Sciences and Engineering Research Council of Canada (RGPIN-2022-04774). T.L. is supported by the Fonds de Recherche du Québec – Santé (FRQS) fellowship (353841). A.A. is supported by the CRBS studentship and McGill Internal studentship. The research reported in this publication was supported by the University of Michigan Cryo-EM Facility (U-M Cryo-EM). U-M Cryo-EM is grateful for support from the U-M Life Sciences Institute and the U-M Biosciences Initiative. We acknowledge the following funding sources: S10OD030275 (N.V. & M.A.C) and R01GM141119 (N.V. and M.A.C.).

## Author contributions

Conceptualization: M.C., K.H.B. Methodology: T.A., A.A., N.V, H.G., N.V., M.C., T.L., K.H.B. Investigation: T.A., A.A., N.V., H.G., H.N.N., N.N.D., M.H.N., T.L., M.C., K.H.B. Visualization: T.A., A.A., H.G., and K.H.B. Funding acquisition: M.C., K.H.B. Supervision: M.H.N., T.L., M.C., K.H.B. Writing – original draft: T.A., A.A., H.G., M.C., K.H.B. Writing – review and editing: T.A., A.A., N.A., M.C., T.L., and K.H.B.

## Disclosure of AI use

The authors declare the use of generative AI in the research and writing process. The following tasks were delegated to GAI tools under full human supervision: - Code generation - Proofreading and editing. The GAI tool used was: ChatGPT-5.0, Gemini 2.5.

## Supplemental Figures

**Supplementary Fig. 1:**
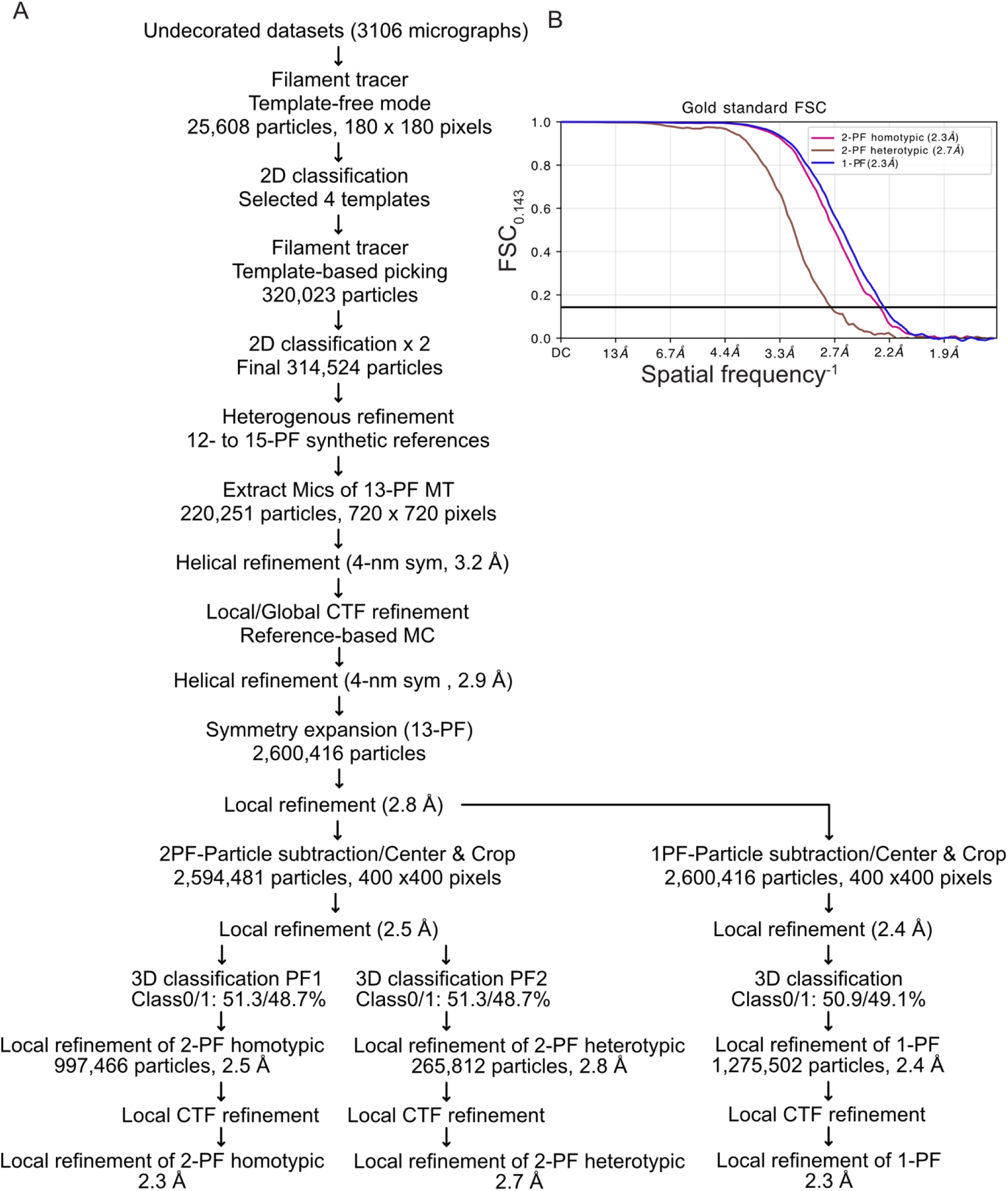
Processing undecorated data of taxol stabilized human MT. (Vangos et al. 2026) (A) Processing workflow of undecorated data. (B) Fourier shell correlation (FSC) of the 2-PF heterotypic, 2-PF homotypic and 1-PF map.

**Supplementary Fig. 2:**
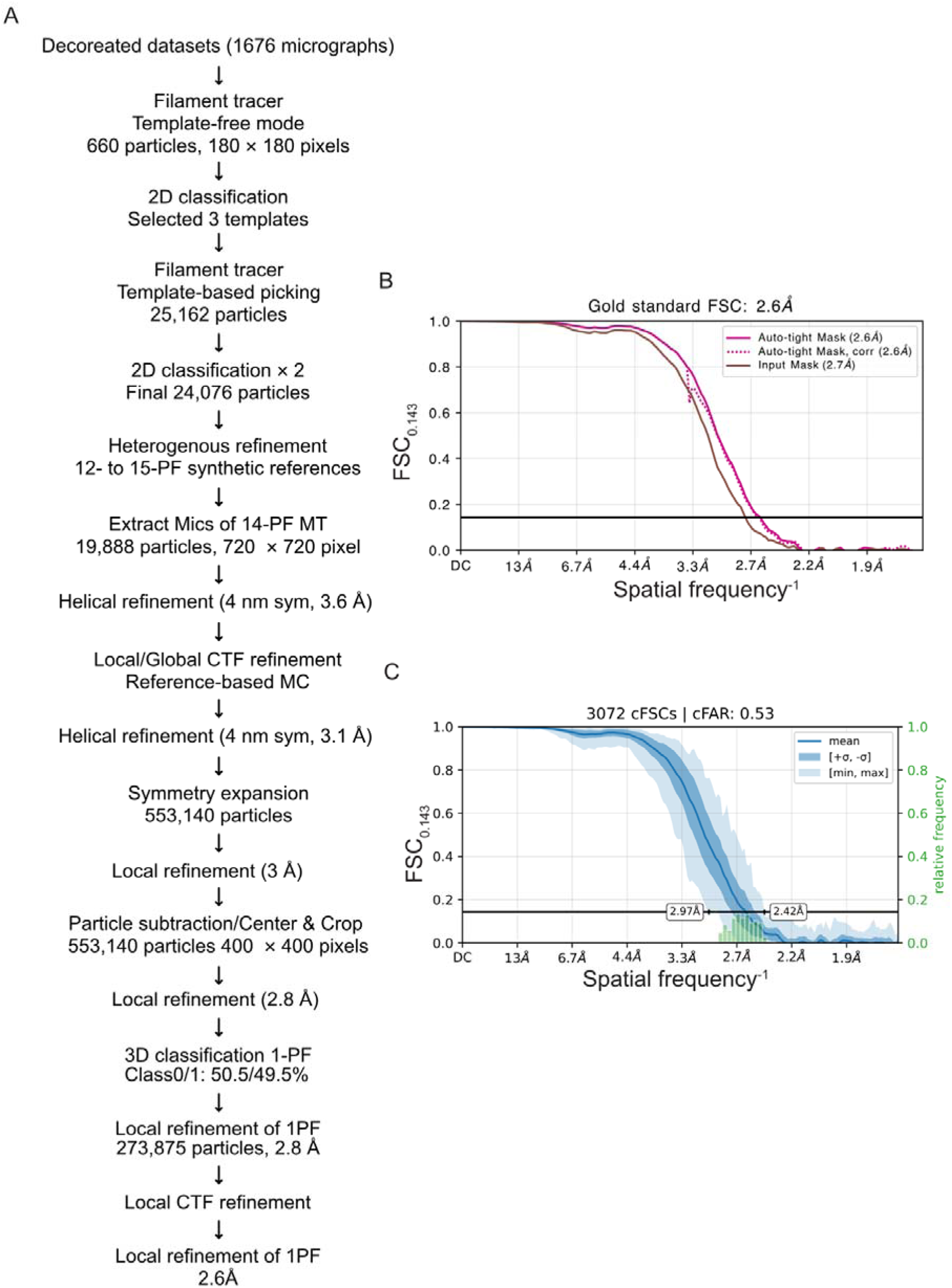
Processing of cryo-EM dataset EMPIAR-13372. (Zhang et al. 2026). (A) Processing workflow of MT decorated with KIF5A (EMPIAR-13372). (B) Fourier shell correlation (FSC) of 1-PF reconstruction obtained in A. (C) Conical Fourier Shell Correlation (cFSC) of the 1-PF reconstruction obtained in A.

## Supplemental Figures

**Supplemental Table S1.** : Cryo-EM data collection and processing

|  | **Undecorated** |  |  | **Decorated** |
| --- | --- | --- | --- | --- |
| **EMPIAR** | (Vangos et al. 2026) |  |  | EMPIAR-13372 |
| **Reconstruction** | 2-PF homotypic | 2-PF heterotypic | 1-PF | 1-PF |
| **Microscope** | Titan Krios G3 |  |  | Titan Krios |
| **Voltage (keV)** | 300 |  |  | 300 |
| **Detector** | K3 BioQuantum |  |  | K2 Summit |
| **Magnification** | 105,000 X |  |  | 78,125 X |
| **GIF slit width** | 20 eV |  |  | 20 eV |
| **Number of movies** | 3,197 |  |  | 1,676 |
| **Number of frames** | 50 |  |  | 62 |
| **Exposure time (s)** | 12 |  |  | 13 |
| **Total dose (e/Å<sup>2</sup>)** | 50 |  |  | 65 |
| **Original pixel size (Å/pixel)** | 0.834 |  |  | 0.64 |
| **Processing pixel size** | 0.834 | 0.834 | 0.834 | 0.835 |
| **Initial number of particles** | 220,251 | 220,251 | 220,251 | 19,888 |
| **Symmetry imposed** | C1 | C1 | C1 | C1 |
| **Final number of particles** | 997,466 | 265,821 | 1,275,502 | 273,875 |
| **Final processing box size** | 400 | 400 | 400 | 400 |
| **Map resolution (Å)** | 2.3 | 2.7 | 2.3 | 2.6 |
| **Model building** | Rigid body fit | Rigid body fit | Rigid body fit | Rigid body fit |
| **Accession number** | EMD-78103 | EMD-78104 | EMD-78112 | EMD-78245 |

**Supplemental Table S2:**
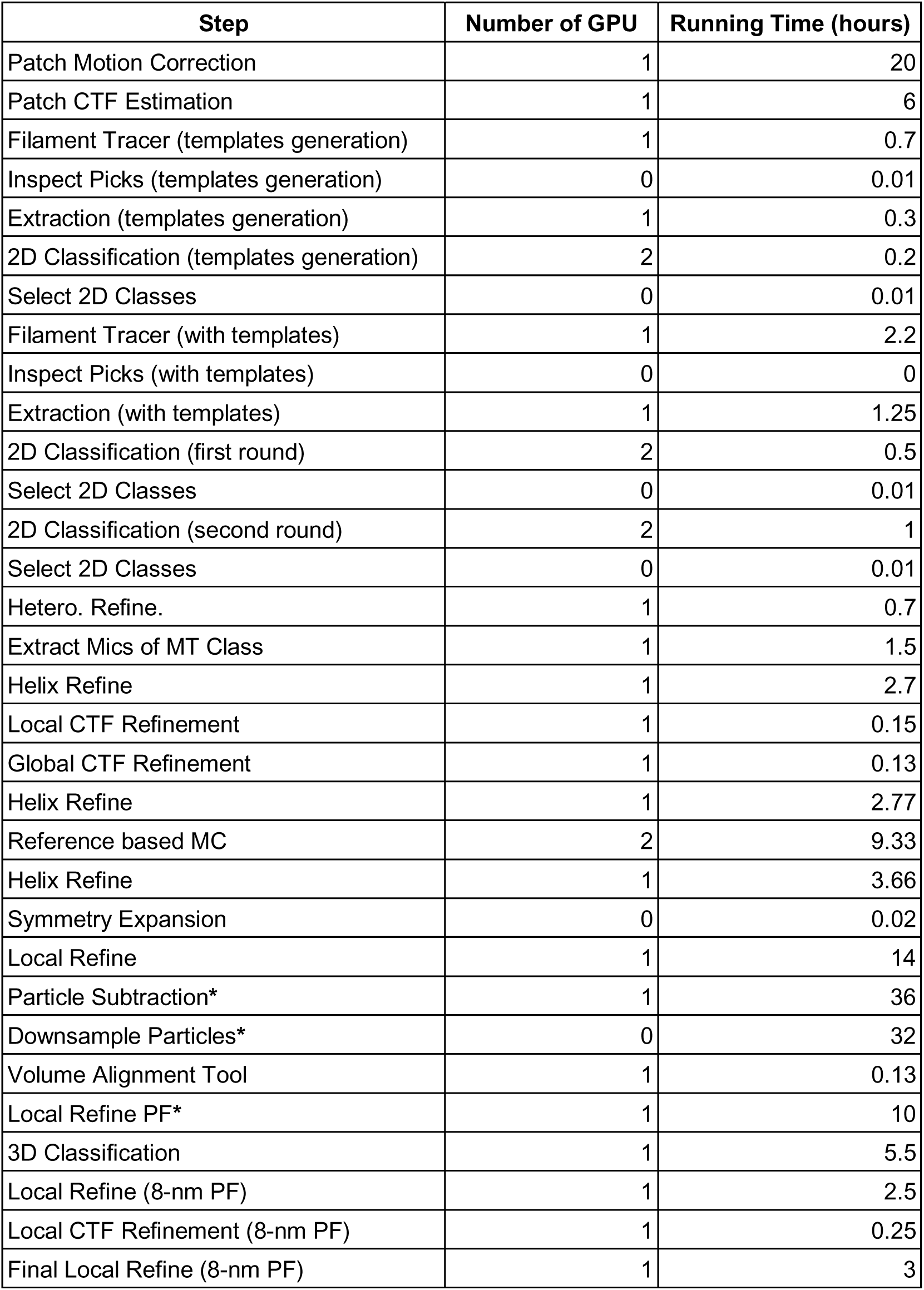
Cryo-EM data processing time for undecorated MT. * denoted steps affected heavily by data read/write speed in our cluster system.

| Step | Number of GPU | Running Time (hours) |
| --- | --- | --- |
| Patch Motion Correction | 1 | 20 |
| Patch CTF Estimation | 1 | 6 |
| Filament Tracer (templates generation) | 1 | 0.7 |
| Inspect Picks (templates generation) | 0 | 0.01 |
| Extraction (templates generation) | 1 | 0.3 |
| 2D Classification (templates generation) | 2 | 0.2 |
| Select 2D Classes | 0 | 0.01 |
| Filament Tracer (with templates) | 1 | 2.2 |
| Inspect Picks (with templates) | 0 | 0 |
| Extraction (with templates) | 1 | 1.25 |
| 2D Classification (first round) | 2 | 0.5 |
| Select 2D Classes | 0 | 0.01 |
| 2D Classification (second round) | 2 | 1 |
| Select 2D Classes | 0 | 0.01 |
| Hetero. Refine. | 1 | 0.7 |
| Extract Mics of MT Class | 1 | 1.5 |
| Helix Refine | 1 | 2.7 |
| Local CTF Refinement | 1 | 0.15 |
| Global CTF Refinement | 1 | 0.13 |
| Helix Refine | 1 | 2.77 |
| Reference based MC | 2 | 9.33 |
| Helix Refine | 1 | 3.66 |
| Symmetry Expansion | 0 | 0.02 |
| Local Refine | 1 | 14 |
| Particle Subtraction* | 1 | 36 |
| Downsample Particles* | 0 | 32 |
| Volume Alignment Tool | 1 | 0.13 |
| Local Refine PF* | 1 | 10 |
| 3D Classification | 1 | 5.5 |
| Local Refine (8-nm PF) | 1 | 2.5 |
| Local CTF Refinement (8-nm PF) | 1 | 0.25 |
| Final Local Refine (8-nm PF) | 1 | 3 |

